# hNav1.5α forms an antiparallel intracellular homodimer but is incorporated into the plasma membrane as a monomer

**DOI:** 10.64898/2026.07.12.738063

**Authors:** Linlu Li, Günther Schmalzing

## Abstract

Electrophysiological studies have long treated the cardiac voltage-gated sodium channel Nav1.5α (SCN5A) as a monomeric pore-forming unit, consistent with all available cryo-EM structures, which show only monomeric architectures. In contrast, biochemical studies — cross-linking, single-molecule pulldown, and native electrophoresis — have reported ∼500 kDa Nav1.5α homodimers with coupled gating. To reconcile this apparent discrepancy, we combined selective labeling of the total (metabolic [^35^S]methionine) and plasma-membrane (membrane-impermeant IRDye 800CW) pools of hNav1.5α with high-resolution clear native electrophoresis (hrCNE) in *Xenopus laevis* oocytes. Total hNav1.5α migrated predominantly as a homodimer that dissociated into monomers upon denaturation, whereas surface-labeled hNav1.5α migrated exclusively as a monomer, confirming mature, Golgi-processed glycosylation by Endo H/PNGase F digestion. This monomer-dimer distribution was unaffected by co-expression with hNavβ1–β4 subunits. Using an orthogonal SpyCatcher/SpyTag covalent tagging strategy, we captured the intracellular homodimer as an irreversible ∼500 kDa complex, and engineered TEV protease cleavage sites revealed that the two protomers associate in a previously unrecognized antiparallel, cyclic arrangement. AlphaFold2-Multimer confidently predicted a monomeric hNav1.5α fold but failed to generate a high-confidence homodimer interface, indicating that this arrangement is not strongly sequence-encoded. Together, our data resolve the electrophysiology-biochemistry discrepancy: hNav1.5α assembles as an antiparallel homodimer in intracellular compartments, likely subject to quality control, but is delivered to the plasma membrane — the physiologically conducting compartment — exclusively as a monomer, irrespective of β-subunit association.

## Introduction

The cardiac voltage-gated sodium channel Nav1.5α, encoded by SCN5A, is essential for the initiation and rapid upstroke of the cardiac action potential, and its dysfunction underlies a broad spectrum of life-threatening arrhythmia syndromes. Nav1.5α exhibits the canonical topology of voltage-gated sodium channel α-subunits, comprising 2,016 amino acid residues organized in four homologous domains (DI–DIV), each containing six transmembrane segments (S1–S6) with associated pore-forming and voltage-sensing elements (Catterall, 2000, 2012). Biochemical and functional evidence indicates that Nav1.5α-subunits can associate as homodimers: Clatot et al. showed by cross-linking and single-molecule pulldown that Nav1.5α forms a complex of >500 kDa with coupled gating, mediated both by direct interaction via the intracellular DI–DII linker and by the adaptor protein 14-3-3 (Clatot et al., 2017; Clatot et al., 2012). High-resolution clear native PAGE (hrCNE) has provided independent biochemical support for Nav1.5α–α interactions (Ruhlmann et al., 2020).

In striking contrast, high-resolution cryo-EM structures of eukaryotic Nav channels, including several structures of human Nav1.5α (hNav1.5α) in different functional and pharmacological states, have so far revealed only monomeric channel architectures, without clear evidence for homodimeric organization (Biswas et al., 2025; Jiang et al., 2021; Li et al., 2021; Pan et al., 2021; Pan et al., 2018; Shen et al., 2017). A notable exception is a crystal structure of the isolated Nav1.5α C-terminal cytoplasmic domain in complex with calmodulin, which revealed an asymmetric Nav1.5α–Nav1.5α contact formed by the EF-hand-like domain and helix αVI, with several arrhythmia-associated mutations mapping to this interface (Gabelli et al., 2014); this structure, however, comprises only the soluble cytoplasmic domain rather than the full-length channel, leaving open whether it reflects the assembly of the intact, membrane-embedded protein.

Therefore, we re-investigated the oligomeric state of hNav1.5α using two approaches: (1) labeling of the total pool of hNav1.5α channels with [^35^S]methionine and, selectively, the plasma-membrane pool with a lysine-reactive, membrane-impermeant fluorescent dye, and (2) a SpyCatcher-based covalent tagging approach (Keeble et al., 2019) (re-proving the formation and orientation of hNav1.5α homodimers. We found that hNav1.5α channels exist in the plasma membrane exclusively as monomers, either alone or in complex with Nav1.5β subunits, whereas intracellular hNav1.5α channels form antiparallel homodimers in *X. laevis* oocytes.

## Materials and Methods

### *X. laevis* care and oocyte preparation

Adult *X. laevis* frogs were maintained and partially, surgically ovariectomized according to procedures approved by the local animal welfare committee (Düsseldorf, Germany; reference no. 8.87-51.05.20.10.131) in compliance with EC Directive 86/609/EEC for animal experiments. Fully collagenase defolliculated Dumont stage V–VI oocytes were prepared as described previously and used for expression experiments (Schmalzing & Markwardt, 2022).

### Plasmid preparation

The oocyte expression construct hNav1.5α^S3^, encoding full-length hNav1.5α with a C-terminal S3 tag (SAWSHPQFEKGGGSGGGSGGSAWSHPQFEK) in the pNKS2 oocyte expression vector (Gloor et al., 1995), was available from our previous study (Ruhlmann et al., 2020). In addition, an hNav1.5α^S4^ construct with two S3 tags fused in tandem with an intervening GGGSGGGSGGS linker was generated to test whether a duplicated S3 tag would increase recovery. Plasmids encoding hNavß2 and hNavß3 subunits were obtained from the Harvard plasmid repository (hSCN2B: pDONR221, HsCD00044489; hSCN3B: pDONR201, HsCD00081587), and plasmids encoding hSCN1B^synth^ and hSCN4B^synth^ (NM_174934.3) were ordered as custom synthetic plasmids from BioCat GmbH (Heidelberg, Germany). All β-subunit coding sequences were subcloned by Gateway LR recombination into a Gateway-compatible version of pNKS2 (pNKS2_GW) and fused to a C-terminal monomeric GFP (mGFP, i.e. ^A206K^GFP; (von Stetten et al., 2012)).

### cRNA synthesis and expression

Plasmids were linearized at a unique XhoI restriction site downstream of the long poly(A) tail, which is crucial for efficient translation in *X. laevis* oocytes (Schmalzing et al., 1992). Capped cRNAs were synthesized using SP6 polymerase, rNTPs and the 5’ anti-reverse cap analog (ARCA) m_2_^7,3’-O^GP_3_G, purified as described previously, and stored at -80 °C until use (Schmalzing & Markwardt, 2022). Defolliculated oocytes were injected with the desired cRNA or mixtures of cRNAs as indicated in the respective experiments

### Metabolic [^35^S]methionine labeling and surface IRDye 800CW labeling

For metabolic labeling, cRNA-injected oocytes were incubated overnight in methionine-free medium supplemented with [^35^S]methionine to pulse-label newly synthesized proteins, followed by a 24 h chase. On day 2 after injection, surface-exposed proteins were labeled with the membrane-impermeant IRDye 800CW reagent under conditions that promote covalent attachment to plasma membrane proteins. This procedure resulted in strong IRDye 800CW signal from surface ^S3^hNav1.5α, whereas associated hNavβ^GFP^ subunits were detected via their intrinsic GFP fluorescence. Under enhanced contrast settings, IRDye 800CW surface labeling of hNavβ^GFP^ subunits also became visible, but only at the expense of a high background fluorescence (data not shown).

### Detergent extraction, Strep-Tactin purification, SDS-PAGE and hrCNE

Oocytes were extracted in digitonin-containing buffer under native conditions to solubilize plasma membrane and intracellular Nav channel complexes, as described previously (Ruhlmann et al., 2020). This approach builds on our earlier work in which digitonin, a particularly mild detergent, combined with BN-PAGE or hrCNE faithfully reported the oligomeric structure of various membrane proteins such as homodimeric anoctamins (Fallah et al., 2011; Stolz et al., 2015), homo- and heterotrimeric P2X receptors (Aschrafi et al., 2004; Becker et al., 2008; Nicke et al., 1998), trimeric acid-sensing ion channels (Fischer et al., 2023) and pentameric glycine receptors (Griffon et al., 1999; Haeger et al., 2010). By analogy, we purified S3-tagged hNav1.5α constructs and associated hNavβ^GFP^ subunits from digitonin extracts using Strep-Tactin Sepharose beads under non-denaturing conditions. Although Ni-NTA is adequate for purifying single recombinant proteins, we used the more specific Strep-tag/Strep-Tactin system here to minimize nonspecific contaminants in native multimeric Nav complexes.

After repeated washing of the respective beads, bound proteins were eluted with desthiobiotin- or imidazole-containing buffer and directly loaded onto self-cast denaturing and native PAGE gels (Wittig et al., 2010; Wittig et al., 2006; Wittig et al., 2007). Immediately after electrophoresis, the wet gels were carefully placed onto the glass surface of the ChemiDoc MP imaging system (Bio-Rad) and scanned for IRDye 800CW plasma-membrane labeling and total hNavβ^GFP^ fluorescence. The gels were then dried on filter paper and subsequently exposed to phosphorimaging screens for several days to detect metabolically incorporated [^35^S]methionine in hNav1.5α^S3^ constructs and co-purified hNavβ^GFP^ subunits using a Storm phosphorimager. Note that co-purified hNavβ^GFP^ fluorescence primarily reports β–α association rather than the subcellular distribution of the β subunits. In the [^35^S] phosphorimager scans, only the hNav1.5α bands are readily visible under the chosen imaging conditions, whereas [^35^S] signals from hNavβ^GFP^ can be detected only after strong contrast enhancement, at which point the hNav1.5α signals become saturated.

### High-resolution clear native electrophoresis (hrCNE)

Native oligomeric states of hNav1.5α were analyzed by hrCNE using native Strep-Tactin eluates. [^35^S]methionine-labeled hNav1.5α species were visualized by phosphorimaging; single and double dot annotations in the figures indicate monomeric and homodimeric forms, respectively.

### Use of AI tools

The AI language model Claude (Claude Sonnet 5, Anthropic) was used to assist in revising and elaborating the manuscript text, and in identifying additional relevant references. The tool was not used for data analysis, interpretation of results, or figure generation. All AI-generated content and suggested references were reviewed and verified by the authors, who take full responsibility for the accuracy and integrity of the final manuscript.

## Results

### Effect of Navß subunits on the oligomeric state of hNav1.5 in the plasma membrane and intracellular compartments

As binding affinity increases with a Twin-Strep-tag® containing two Strep-tagII motifs in tandem separated by a flexible GGGSGGGSGGS linker, referred to as an S3-tag (Junttila et al., 2005; Schmidt et al., 2013), we generated an extended variant designated S4-tag by duplicating the S3-tag; S4 was fused N-terminally (^S4^hNav1.5α), whereas S3 was fused C-terminally (hNav1.5α^S3^). Following metabolic [^35^S]methionine labeling combined with StrepTactin purification, we observed similar recovery from *X. laevis* oocytes with both tags and predominant migration as homodimers the hrCNE gel, which dissociated into monomers upon LiDS treatment (Fig. 1A). The somewhat faster migration of ^S4^hNav1.5α and hNav1.5α^S3^ monomers in LiDS-treated samples can be attributed to a charge effect caused by the denaturing LiDS.

**Fig. 1.**
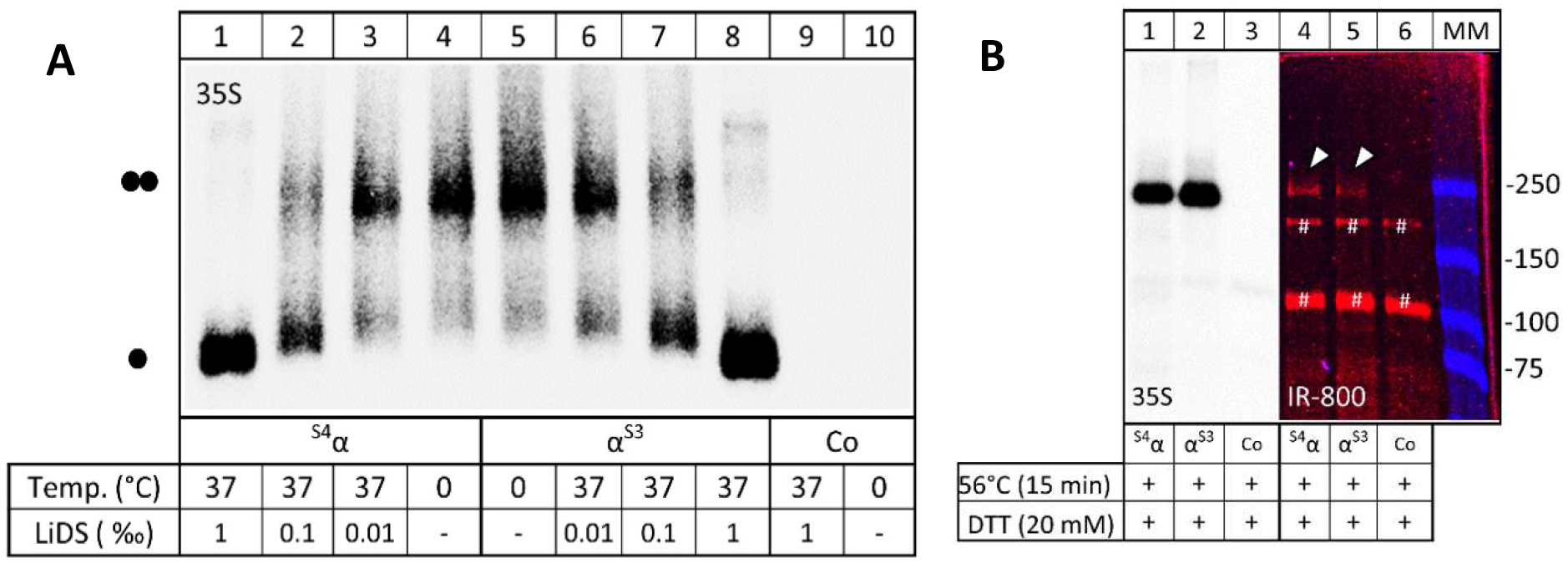
Comparison of the efficiency of purifying hNav1.5α via an S3 or S4 tag. *X. laevis* oocytes expressing the indicated constructs were pulse-labeled overnight with [^35^S]methionine and surface-labeled at day 2 post-injection with the membrane-impermeant fluorescent dye IRDye 800CW. Detergent extracts (1% digitonin in 0.1 M Na-phosphate buffer, pH 8.0) were purified under non-denaturing conditions using Strep-Tactin Sepharose. Non-injected oocytes (Co, control) were processed in parallel. (**A**) hrCNE. Samples were incubated with increasing LiDS concentrations to achieve progressive denaturation, resolved by hrCNE, and scanned for incorporated [^35^S]methionine using a Storm phosphorimager. Note that increasing LiDS concentrations resulted in progressive dissociation of homodimers (••) into monomers (•). (**B**) Aliquots of the same samples were denatured in SDS sample buffer, resolved by SDS-PAGE, and analyzed for total [^35^S]-methionine incorporation (intracellular plus plasma membrane) and plasma-membrane-bound IR800 fluorescence using [^35^S]phosphorimaging and a ChemiDoc MP imaging system, respectively. Arrows indicate plasma membrane-bound hNav1.5α, absent in the control (lanes 3 and 6), while the hash symbol (#) marks co-purified Cy5-labeled background proteins present in all three samples. Together, these data demonstrate that both S3- and S4-tagged constructs permit efficient recovery of hNav1.5α from native digitonin extracts with comparable yield and purity, and we therefore used smaller the S3-tag as the standard configuration for subsequent experiments.

Cell-surface IRDye 800CW labeling of the same samples as in Fig. 1A likewise revealed no evident difference in the comparably weak plasma membrane expression between ^S4^hNav1.5α and hNav1.5α^S3^ (Fig. 1B). Together with the similar [^35^S]methionine-based recovery of both constructs after StrepTactin purification (Fig. 1A), this indicates that the S3-tag already enables efficient recovery of hNav1.5α.

Aliquots of the natively eluted samples were incubated with increasing LiDS concentrations for 1 h at 37 °C to induce graded denaturation (Fig. 1A), and ion channel proteins were subsequently separated by hrCNE and visualized by [^35^S]phosphorimaging. A single dot (•) marks the hNav1.5α monomer, whereas two dots (••) indicate the hNav1.5α homodimer. Taken together, both tags support comparable purification and detection of hNav1.5α of homodimeric and monomeric hNav1.5α, and we therefore chose the smaller S3 tag for all subsequent experiments.

### Role of hNavß subunits in hNav1.5α surface expression

Navβ subunits have been reported to modulate the trafficking and membrane localization of Nav1.5α (Brackenbury & Isom, 2011; Hull & Isom, 2018). To examine whether hNavβ subunits affect plasma membrane and/or total expression of hNav1.5α in *X. laevis* oocytes, we co-expressed ^S3^hNav1.5α with hNavβ3^GFP^ over a wide range of α:β1 ratios (from 1:1 to 1:30; Fig. 2A) and, in separate experiments, with hNavβ2^GFP^ or hNavβ3^GFP^ at a fixed α:β cRNA ratio of 1:2 (Fig. 2B). To specifically visualize the plasma membrane fraction of ^S3^hNav1.5α in these co-expression experiments, we applied the lysine-reactive, membrane-impermeant IRDye® 800CW to intact oocytes, restricting fluorescence to the extracellularly accessible surface pool. The resulting IRDye-labeled Strep-Tactin eluates were first analyzed by SDS-PAGE to quantify surface versus total hNav1.5α expression (Fig. 2A,B) and, in subsequent experiments, by BN-PAGE under native and LiDS-denaturing conditions to assess the oligomeric state of the same preparations (Fig. 3).

**Fig. 2.**
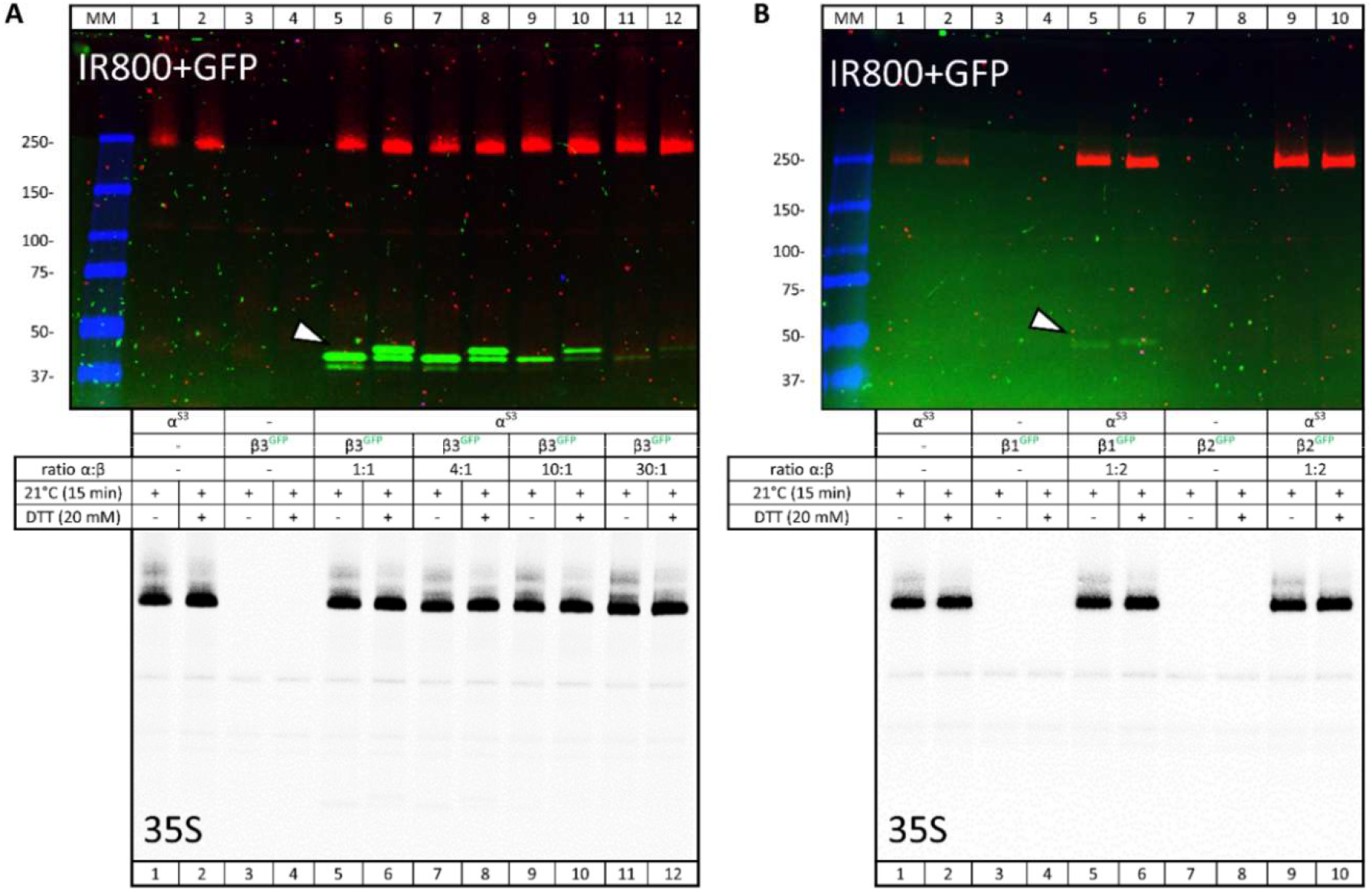
Co-purification of hNavβ subunits with surface hNav1.5α. hNav1.5α^S3^ was expressed as indicated, either alone or together with hNavβ1^GFP^, hNavβ2^GFP^, or hNavβ3^GFP^, by cRNA co-injection into *X. laevis* oocytes. Oocytes were pulse-labeled overnight with [^35^S]methionine followed by a 24 h chase. On day 2, surface proteins were labeled with the membrane-impermeant dye IRDye 800CW. Proteins were extracted with 1% digitonin, purified on Strep-Tactin–Sepharose under non-denaturing conditions, and then denatured with 0.2% SDS under non-reducing or reducing conditions (20 mM DTT). Aliquots were resolved by SDS-urea-PAGE and visualized by fluorescence scanning (ChemiDoc MP; **A,B**) and ([^35^S]phosphorimaging (Storm; **C,D)**. (A,B) Fluorescence scan detecting IRDye 800CW-labeled plasma-membrane-bound hNav1.5α^S3^ as ∼250 kDa bands and co-purified β-GFP subunits as ∼40 kDa bands in the GFP channel. White arrows indicate fully co-purified β-GFP subunits lacking an affinity tag. Note that contrast settings at the two emission wavelengths were adjusted independently to optimize visualization of the respective protein bands; consequently, signal intensities of the green and red channels do not reflect the relative stoichiometry of the two subunits. (**C,D**) [^35^S]Phosphorimager scan of the same gels. The [^35^S]methionine-labeled hNav1.5α subunits are readily detectable as ∼250 kDa bands in all lanes in which hNav1.5α^S3^ is expressed, whereas [^35^S]-labeled hNavβ^GFP^ signals are not visible under the phosphorimager settings used here but become apparent after contrast enhancement (not shown).

**Fig. 3.**
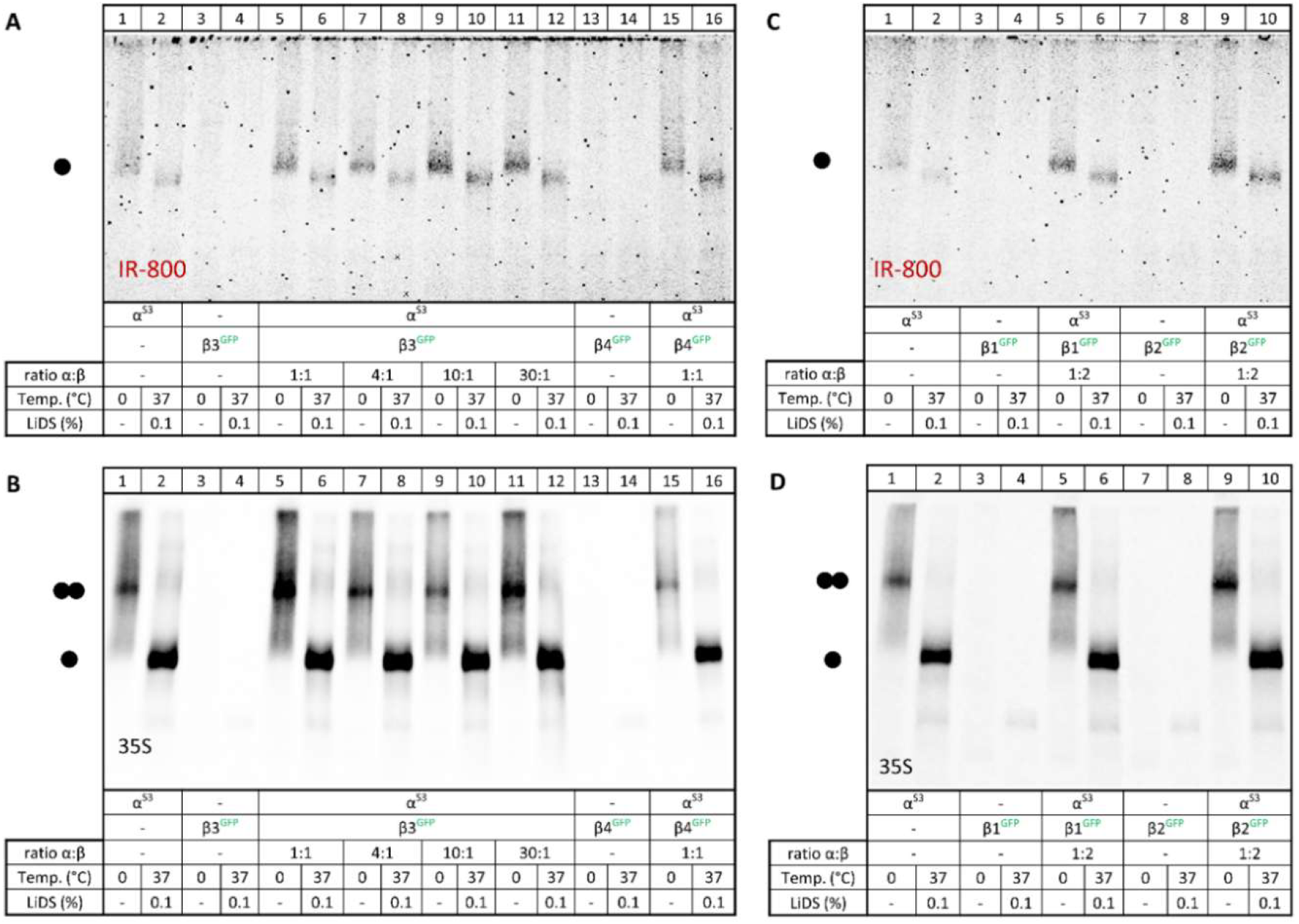
Comparative analysis of the native assembly of plasma membrane–resident hNav1.5α^S3^ and total hNav1.5α^S3^. Protein samples previously used for Fig. 2 were either kept on ice (0°C) or treated with denaturing LiDS at the indicated concentrations in BN-PAGE sample buffer for 1 h at 37°C. Proteins were separated by BN-PAGE and visualized by IR800 ChemiDoc MP fluorescence scanning to assess plasma membrane expression (**A,C**) and Storm [^35^S]phosphorimaging for total expression (**B,D**). A single black dot at the left margin marks the migration position of the hNav1.5α^S3^ monomer; a double dot marks the hNav1.5α^S3^ homodimer. (**A,B**) hNav1.5α^Strep3^, hNavβ3^GFP^ and hNavβ4^GFP^ were expressed either alone or in the indicated combinations. (**A**) IR800-labeled plasma membrane expression; (**B**) [^35^S]methionine-labeled total expression. (**C,D**) hNav1.5α^S3^ was expressed alone or co-expressed with hNavβ1^GFP^ or hNavβ2^GFP^.

For Fig. 2A,B, Strep-Tactin–purified ^S3^hNav1.5α complexes were evaluated by IRDye® 800CW fluorescence (covalent surface labeling of ^S3^hNav1.5α), co-purified hNavβ^GFP^ fluorescence, and metabolic [^35^S]methionine incorporation to assess plasma-membrane abundance and total synthesis of hNav1.5α. Note that co-purified hNavβ^GFP^ fluorescence primarily reports β–α association rather than the subcellular distribution of the β subunits. In the [^35^S] phosphorimager scans, only the hNav1.5α bands are readily visible under the chosen imaging conditions, whereas [^35^S] signals from hNavβGFP can be detected only after strong contrast enhancement, at which point the hNav1.5α signals become saturated. Across these conditions, the hNav1.5α bands labeled with surface IRDye 800CW and with [^35^S]methionine remained comparably in intensity, indicating that changes in β-subunit abundance or subtype have little, if any, impact on plasma-membrane abundance or total synthesis of hNav1.5α under these conditions.

The presence of hNavβ^GFP^ subunits in the fluorescence scans of the Strep-Tactin eluates in Fig. 2A and 2B, despite the absence of an affinity tag on hNavβ^GFP^, indicates that hNav1.5α^S3^ is co-purified in a relatively stable complex with hNavβ^GFP^. hNavβ3^GFP^ remains detectable even when its cRNA is diluted 30-fold relative to hNav1.5α, whereas co-recovery of hNavβ1^GFP^ is clearly weaker and no co-purification of hNavβ2^GFP^ is detectable under the same conditions. This robust co-recovery of hNavβ3^GFP^ with hNav1.5α^S3^ is consistent with previous reports identifying the β3 subunit as an important structural and functional regulator of Nav1.5α, with pronounced effects on Nav1.5α gating, clustering and mechanosensitivity (Salvage et al., 2020).

A direct, yet weak interaction between hNav1.5α and hNavβ1 is also evident, as hNavβ1^GFP^ co-purified upon co-expression with hNav1.5α^S3^ (Fig. 2B, upper scan, lanes 5 and 6; hNavβ1^GFP^ indicated by a white arrow), consistent with previous reports on direct Nav1.5α-ß1 association (Brackenbury & Isom, 2011; Hull & Isom, 2018). In contrast to hNavβ1 and hNavβ3, no interaction of hNavβ2 with hNav1.5α, either covalent or non-covalent, was detected (Fig. 2B, upper scan, lanes 9 and 10). If hNavβ2^GFP^ were linked to hNav1.5α^S3^ via disulfide bonds, as reported for some other Nav isoforms (Brackenbury & Isom, 2011; Hull & Isom, 2018), an SDS-resistant protein band corresponding to the sum of their molecular masses (271 kDa: hNav1.5α^S3^ 220 kDa + hNavβ1^GFP^ 51 kDa) and migrating more slowly than hNav1.5α^S3^ alone would be expected, but no slower migrating band is visible in Fig. 2B. A non-covalent association between the two subunits is therefore the most plausible explanation for the co-isolation of hNavβ1^GFP^ and hNavβ3^GFP^ subunits, which is lost under the denaturing conditions of the SDS-urea-PAGE in Fig. 2.

## Identification of hNav1.5α as monomeric in the plasma membrane and homodimeric in intracellular membranes

The IRDye® 800CW-labeled plasma membrane pool and the [^35^S]-labeled total hNav1.5α pool defined in Fig. 2 were further analyzed by high-resolution clear native electrophoresis (hrCNE) to assess the oligomeric state of the same Strep-Tactin eluates (Fig. 3). We found that the IR800-labeled surface hNav1.5α migrated entirely as a monomer, whereas the [^35^S]-metabolically labeled hNav1.5α appeared predominantly as a homodimer in the same hrCNE gel, thereby identifying the monomer as the plasma membrane-expressed species (Fig. 3A,C). The [^35^S]-labeled fraction, representing predominantly intracellular hNav1.5α, migrated as homodimer that dissociated into its monomers upon denaturing treatment with 0.1% LiDS prior to hrCNE analysis (Fig. 3B,D).

Taken together, our biochemical analysis demonstrates that, despite the presence of homodimeric hNav1.5α assemblies under native extraction conditions (Ruhlmann et al., 2020) (Fig. 3B,D), these homodimers may represent an exclusively intracellular pool of membrane-inserted channels residing in early biosynthetic or other internal membrane compartments. This interpretation is consistent with intracellular quality control mechanisms acting on non-productive oligomeric assemblies in internal membrane compartments under heterologous expression conditions. Moreover, the biochemical detection of mature hNav1.5α as a monomeric channel in the plasma membrane (Fig. 3A,C) also agrees with the classical electrophysiological view of Nav1.5 as a monomeric pore-forming unit in functional studies (Morris & Juranka, 2007).

To further substantiate the plasma membrane localization of IR800-labeled monomeric hNav1.5α, we performed N-glycan analysis using Endo H and PNGase F (Fig. 4). SDS-PAGE followed by IR800 fluorescence scanning revealed Endo H-resistant but PNGase F-sensitive plasma membrane–associated hNav1.5α, consistent with complex, Golgi-processed N-glycans as expected for mature, plasma membrane-localized hNav1.5α channels (Fig. 4A,C). In parallel, we analyzed the [^35^S]methionine-labeled hNav1.5α pool using the same Endo H/PNGase F digestion strategy. In contrast to the plasma membrane fraction, this metabolically labeled pool was sensitive to both Endo H and PNGase F (Fig. 4B,D), indicating immature, high-mannose N-glycans and suggesting that a substantial fraction of hNav1.5α resides in the endoplasmic reticulum and early secretory pathway. Accordingly, the Endo H-resistant, PNGase F-sensitive species corresponds to properly Golgi-processed, plasma membrane–localized hNav1.5α channels are present as monomers in the plasma membrane.

**Fig. 4.**
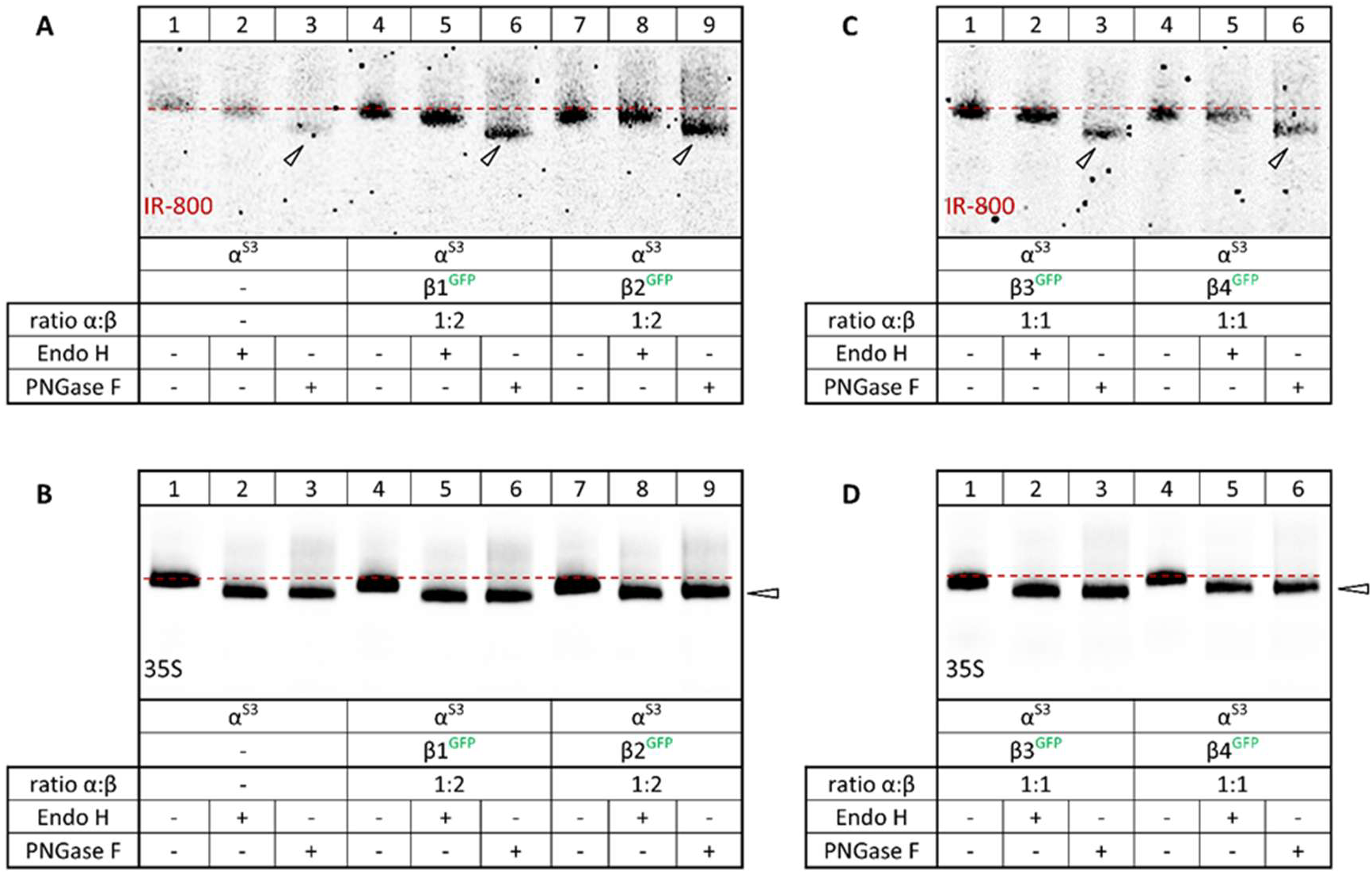
Analysis of the N-glycosylation status of total and plasma membrane hNav1.5α^S3^. Selected samples from Fig. 3 were denatured with 0.2% SDS and 20 mM DTT and incubated with Endo H or PNGase F as indicated, then analyzed by urea–SDS-PAGE and visualized by Storm [^35^S]phosphorimaging and ChemiDoc MP fluorescence scanning. (**A,C**) Plasma membrane labeled IR800-hNav1.5α expressed alone or together with any of the three hNavβ1 subunits was Endo H resistant, but PNGase sensitive, consistent with plasma membrane location. (**B,D**) In contrast, metabolically labeled ^35^S-hNav1.5α expressed alone or together with any of the three hNavβ1 subunits was Endo H and PNGase sensitive, consistent with location in intracellular compartments.

### 4.1 SpyCatcher/SpyTag approach to the homodimerization of hNav1.5^**α**^

To probe the oligomeric state of hNav1.5α with an orthogonal strategy, we employed the SpyCatcher/SpyTag (SpyC/SpyT) system, which mediates irreversible isopeptide bond formation between a short peptide tag (SpyTag) and its cognate partner (SpyCatcher) (Zakeri et al., 2012). We used the optimized variants SpyTag003 (16 residues) and SpyCatcher003 (145 residues), which display improved stability, solubility and reaction kinetics (Keeble & Howarth, 2020), without explicitly denoting the “003” suffix in construct names.

Fig. 5 shows that hNav1.5α^mGFP^ and ^SpyC^hNav1.5α^mGFP^ migrated at identical positions (∼250 kDa), indicating that the SpyC tag does not alter the electrophoretic mobility of hNav1.5α. Co-expression of ^SpyC^hNav1.5α^mGFP^ with ^SpyT^hNav1.5α or hNav1.5α^SpyT^ resulted in the appearance of an additional high-molecular-mass band at ∼500 kDa, consistent with the calculated mass of a covalent ^SpyC^hNav1.5α^mGFP^– ^SpyT^hNav1.5α or ^SpyC^hNav1.5α^mGFP^–hNav1.5α^SpyT^ homodimer, respectively (Fig. 5, lanes 5–10). Importantly, this homodimer band persisted under denaturing SDS–PAGE conditions, demonstrating that the two subunits are covalently linked via SpyC–SpyT isopeptide bond formation rather than held together by non-covalent interactions. The relatively weak fluorescence intensity of the dimer band compared to the monomer reflects the fact that only one subunit of each homodimer carries mGFP.

**Fig. 5.**
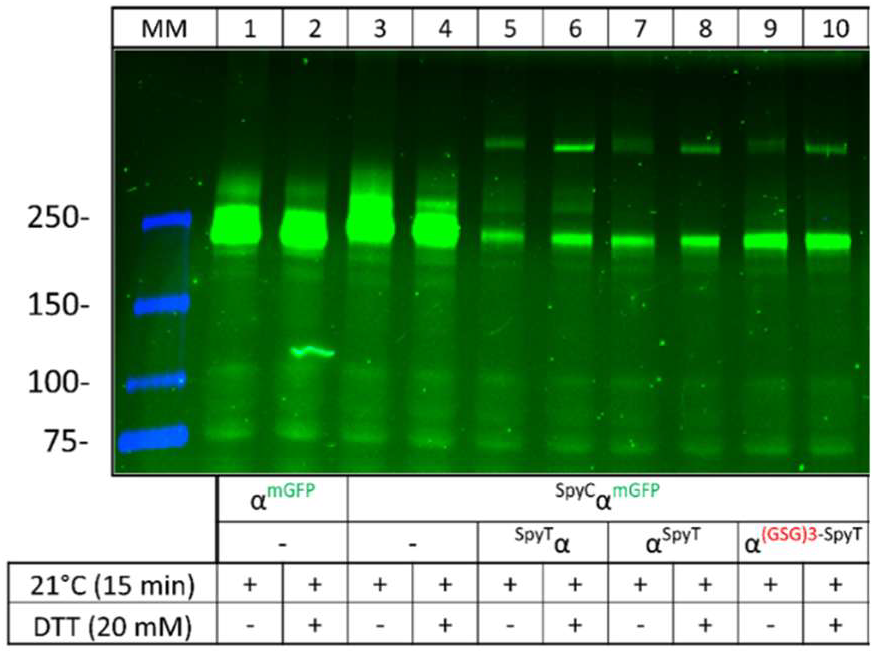
SpyC–SpyT tagging results in covalent hNav1.5α homodimerization. In construct names, SpyC or SpyT is placed N-terminally (^SpyC^hNav1.5α^mGFP, SpyT^hNav1.5α) or C-terminally (hNav1.5α^SpyT^, hNav1.5α^(GSG)^_3_^SpyT)^ relative to hNav1.5α to indicate their respective positions. hNav1.5α^mGFP^ (lanes 1–2) and ^SpyC^hNav1.5α^mGFP^ (lanes 3–4) migrated at identical positions (∼250 kDa according to mass markers). Co-expression of ^SpyC^hNav1.5α^mGFP^ with hNav1.5α carrying an N- or C-terminal SpyT tag resulted in the appearance of an additional high-molecular-mass band that, based on extrapolation from the 150 kDa and 250 kDa markers, yielded 221 kDa and 396 kDa, compared with calculated 232 kDa and 496 kDa, respectively. The deviation in the calculated mass of the dimer likely reflects extrapolation beyond the calibration mass range and the use of soluble protein markers to estimate the mass of a membrane protein.

### 1.5 SpyTag/SpyCatcher-based fluorescence labeling of hNav1.5^**α**^

As an alternative strategy, we fused the SpyTag either to the N-terminus or the C-terminus of hNav1.5α (^SpyT^hNav1.5α and hNav1.5α^SpyT^), while SpyCatcher was fused to monomeric GFP (^SpyC^mGFP). This strategy allows independent expression and folding of hNav1.5α and mGFP by restricting modifications to short peptide tags on hNav1.5α and mGFP instead of full-length GFP fusions. In addition, a SpyCatcher–hNav1.5α^mGFP^ fusion (^SpyC^hNαv1.5α^mGFP_SpyT^) was generated to assess covalent homodimerization via intracomplex SpyC– SpyT ligation between co-expressed subunits.

hNav1.5α variants were expressed without or with ^SpyC^mGFP in *X. laevis* oocytes, and digitonin extracts were analyzed by urea SDS–PAGE (Fig. 6A). Co-expression of ^SpyC^mGFP rendered ^SpyT^hNav1.5α (lanes 7– 8) and hNav1.5α^SpyT^ (lanes 9–10) mGFP-fluorescent via their compatible SpyT tag, with both migrating at the same position as a directly fused hNav1.5α^mGFP^ control (lanes 1–2), demonstrating efficient covalent labeling of hNav1.5α via SpyC/SpyTag ligation regardless of whether SpyTag was placed at the N- or C-terminus. As expected, the hNav1.5α variant carrying only a C-terminal SpyCatcher (hNav1.5α^SpyC^) did not react with SpyC^mGFP^ and remained undetectable in fluorescence scans (Fig. 6A, lanes 5-6). In contrast, despite carrying a C-terminal SpyT tag, the double-tagged ^SpyC^hNav1.5α^SpyT^ was only minimally labeled by ^SpyC^mGFP (lanes 11–12).

**Fig. 6.**
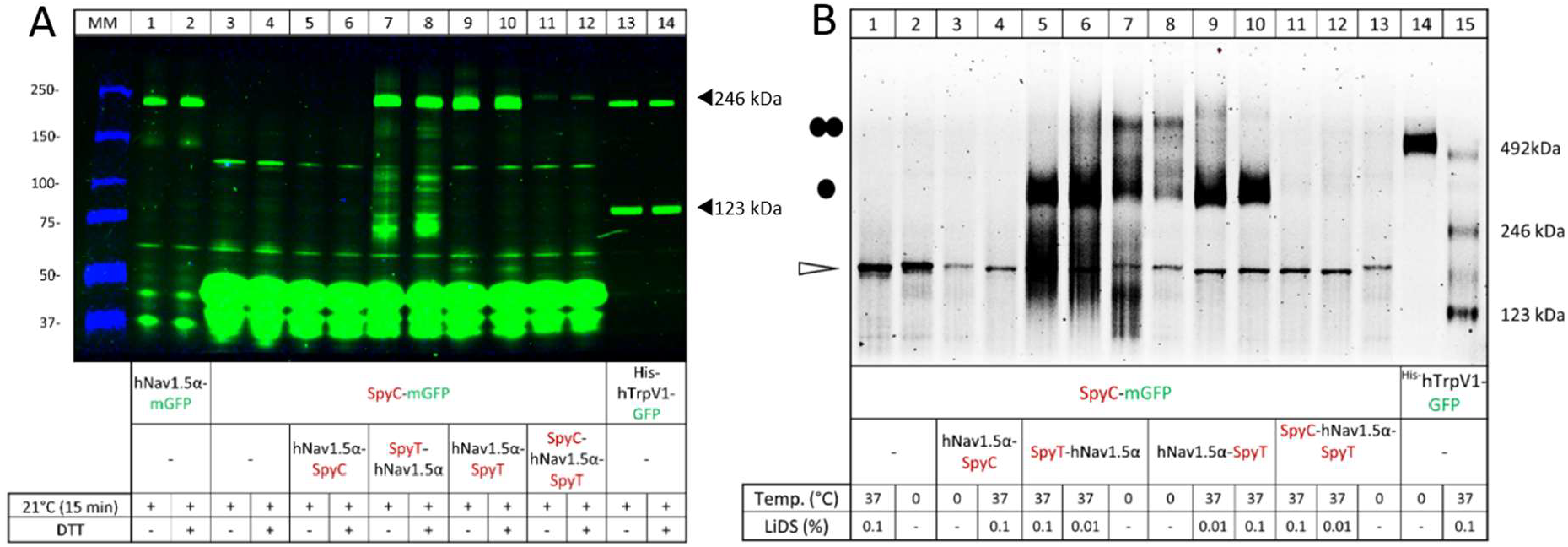
mGFP labeling of hNav1.5α via SpyC/SpyT-mediated protein ligation. The indicated constructs were expressed either alone or in combination in *X. laevis* oocytes, and digitonin extracts were prepared on day 2. In this experiment, hNav1.5α constructs are detectable only if they present a free SpyT tag accessible for covalent conjugation with ^SpyC^mGFP. (**A**) Samples were denatured with 0.2% SDS at 21 °C (-/+ DTT), resolved by 4–10% urea SDS-PAGE, and proteins were visualized by GFP fluorescence scanning. SpyC/SpyT ligation resulted in covalent A labeling of hNav1.5α constructs carrying a free N- or C-terminal SpyT with mGFP, rendering them mGFP-fluorescent, whereas the SpyC-only construct hNav1.5α^SpyC^, which cannot react with ^SpyC^mGFP, remained undetectable. In the N-terminal ^SpyT^hNav1.5α construct, additional lower molecular mass bands may reflect incompletely translated polypeptides that still carry the N-terminal SpyTag and therefore can react with ^SpyC^mGFP. (**B**) hrCNE analysis of hNav1.5α constructs post-translationally labeled with ^SpyC^mGFP. Non-denatured ^SpyT^hNav1.5α covalently labeled with ^SpyC^mGFP is visible as both monomer (•) and homodimer (••). The calculated monomer mass of 310 kDa (^SpyT^hNav1.5α: 256 kDa + ^SpyC^mGFP: 44 kDa) and homodimer mass of 620 kDa match the migration of ^His^Trp^mGFP^-derived mass markers. In contrast, ^SpyC^hNav1.5α^SpyT^ shows only marginal fluorescence upon co-expression of ^SpyC^mGFP, despite carrying a compatible SpyC tag; this is consistent with efficient intracomplex ligation between the N-terminal SpyC and C-terminal SpyT tags, forming an antiparallel covalent homodimer that almost completely prevents intermolecular reaction with ^SpyC^mGFP.

Under native conditions (hrCNE, Fig. 6B), ^SpyT^hNav1.5α and hNav1.5α^SpyT^ visualized via co-expressed ^SpyC^mGFP (lanes 7–8) migrated as ∼500 kDa homodimers, which entirely dissociated into monomers upon incubation with 0.1% LiDS (lanes 5-6 and 9-10). Thus, hNav1.5α labeled via the SpyC/SpyTag system reproduces the homodimeric organization previously observed for directly fused hNav1.5α^mGFP^ (see Fig. 6), while retaining the advantage of a much smaller tag on the hNav1.5α polypeptide. ^SpyC^hNav1.5α^SpyT^ was undetectable under these native conditions (Fig. 6B, lanes 11–13). The virtual inaccessibility of ^SpyC^hNav1.5α^SpyT^ to ^SpyC^mGFP suggests that rapid antiparallel covalent SpyC–SpyT homodimerization of ^SpyC^hNav1.5α^SpyT^ subunits during or shortly after synthesis consumes the available SpyT sites and prevents covalent labeling by ^SpyC^mGFP. Consistent with this interpretation, subsequent experiments showed that ^SpyC^hNav1.5α^SpyT^ preferentially forms antiparallel homodimers via SpyC/SpyTag-mediated ligation (section 3.2.4). which will be addressed later.

Co-expression of ^SpyC^mGFP with ^SpyT^hNav1.5α or hNav1.5α^SpyT^ resulted in additional GFP-positive bands at the migration position of a directly fused hNav1.5α^mGFP^ control, demonstrating efficient covalent labeling of hNav1.5α via SpyC/SpyTag ligation, irrespective of whether SpyTag was placed at the N- or C-terminus (Fig. 6A, lanes 7-10). By contrast, the double-tagged ^SpyC^hNav1.5α^SpyT^ construct (Fig. 6A, lanes 11-12) produced only a weak GFP-positive band, even though a high-molecular-mass ligation product was expected. This behavior is most readily explained by preferential intramolecular or inter-chain SpyC/SpyTag ligation within or between ^SpyT^hNav1.5α^SpyT^ molecules, which would rapidly generate non-fluorescent products invisible in GFP scans and preclude efficient ligation with ^SpyC^mGFP. Consistent with this interpretation, subsequent experiments showed that ^SpyC^hNav1.5α^SpyT^ preferentially forms homodimers via SpyC/SpyTag-mediated ligation (see Fig. 7). As expected, the variant hNav1.5α variant carrying only a C-terminal SpyCatcher (hNav1.5α^SpyC^) did not react with ^SpyC^mGFP and remained invisible in fluorescence scans (Fig. 6A, lanes 11-12).

**Fig. 7.**
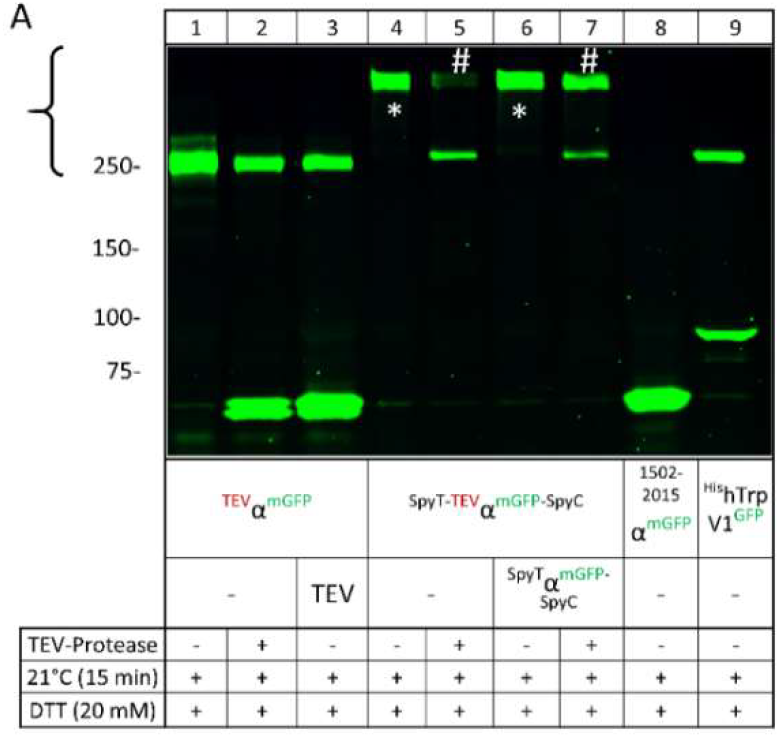
TEV-assisted topology analysis of SpyC–SpyT-linked hNav1.5α dimers. ^1508_TEV^hNav1.5α^mGFP^ and ^SpyT^hNav1.5α^mGFP–SpyC^ were expressed in *X. laevis* oocytes as indicated and analyzed by SDS–PAGE and fluorescence scanning following TEV protease treatment where indicated. **Lane 1**: ^1508_TEV^hNav1.5α^mGFP^ analyzed in the absence of TEV remained uncleaved. **Lanes 2–3**: ^1508_TEV^hNav1.5α^mGFP^ was partially cleaved at the engineered TEV site by co-expressed TEV during expression (lane 2) or by recombinant TEV added to post-expression digitonin extracts (**lane 3**), generating an ∼85 kDa (calculated: 84.4 kDa, unglycosylated) C-terminal mGFP-labeled fragment and a non-visible ∼170 kDa (calculated: 170.0 kDa, unglycosylated) fragment, while the residual uncleaved parental ∼250 kDa band remained detectable. **Lane 4**: Co-expression of ^SpyT-TEV^hNav1.5α^mGFP-SpyC^ resulted in the appearance of a high-molecular-mass ligation product, consistent with a covalent ^SpyT-TEV^hNav1.5α^mGFP-SpyC^ homodimer of 500 kDa, **Lane 5**: Co-expressed TEV protease almost completely converted the SpyT–SpyC-linked homodimers to the ∼250 kDa monomer via the engineered TEV site. **Lane 6**: Co-expression of ^SpyT-TEV^hNav1.5α^mGFP-SpyC^ with ^SpyT^hNav1.5α^mGFP-SpyC^, equal to lane 4, but lacking an TEV-site, resulted in mixed concatamers with 0-2 TEV clevage sites, **Lane 7**: Co-expression of ^SpyT-TEV^hNav1.5α^mGFP-SpyC^ with a ^SpyT–SpyC^-linked hNav1.5α^mGFP^ construct lacking the TEV site produced a mixture of TEV-cleavable and non-cleavable dimers, resulting in only partial TEV-mediated conversion to the ∼250 kDa monomer and a detectable residual dimer band, in contrast to the almost complete monomerization seen in lane 5; lane 8: designed hNav1.5α fragment that encodes exactly the 85 kDa product generated by TEV cleavage ^SpyT-TEV^hNav1.5α^mGFP-SpyC^.

High-resolution clear native electrophoresis (hrCNE) of SpyC/SpyTag-labeled hNav1.5α revealed that ^SpyT^hNav1.5α^SpyC_mGFP^ and hNav1.5α^SpyT_SpyC_mGFP^ complexes migrate as apparent homodimers with an estimated mass of ∼550 kDa, slightly above a ^His^TrpV1^GFP^ tetramer marker (Fig. 6B). Upon denaturation with LiDS at 37°C, these dimers dissociated into monomeric complexes that migrated at the position expected for a single hNav1.5α^mGFP^ equivalent. Thus, hNav1.5α labeled via the SpyC/SpyTag system reproduces the homodimeric organization previously observed for directly fused hNav1.5α^mGFP^, while retaining the advantage of a much smaller tag on the Nav1.5α polypeptide.

### 1.6 TEV-based validation of intracellular hNav1.5α as an antiparallel cyclic homodimer

The refractoriness of ^SpyC^hNav1.5α^SpyT^ to labeling with ^SpyC^mGFP (Fig. 6A, lanes 11-12) is consistent with an antiparallel homodimeric arrangement in which the N-terminal SpyC of one subunit is brought into close proximity to the C-terminal SpyT of the neighboring subunit, enabling inter-subunit covalent self-ligation.

To test this antiparallel cyclic dimer model, we first generated a control by inserting a Tobacco Etch Virus (TEV) protease site at position 1508 of the 2015-residue-long hNav1.5α to generate ^SpyT_1508.TEV^hNav1.5α^mGFP_SpyC^, expecting TEV cleavage to yield two fragments indicative of a monomer: a non-visible, non-GFP–labeled N-terminal fragment of ∼170 kDa (calculated: 170.0 kDa, unglycosylated) and an mGFP-visible C-terminal fragment of ∼85 kDa (calculated: 84.4 kDa, unglycosylated). This is exactly what we observed when we exposed ^1508_TEV^hNav1.5α^mGFP^ to TEV, either by co-expression in *X. laevis* oocytes or by adding recombinant TEV to digitonin extracts of these oocytes (Fig. 7, lanes 1-3). We found that ^1508_TEV^hNav1.5α^mGFP^ migrated as a ∼250 kDa band (lanes 1-3), and, upon TEV treatment by either cRNA co-expression or recombinant protease in digitonin extracts, yielded the predicted ∼85 kDa (calculated: 84.4 kDa, unglycosylated) C-terminal mGFP-labeled fragment, confirming efficient cleavage by both delivery modes (Fig. 7, lanes 2-3). ^1508-2015^hNav1.5α^mGFP^ co-migrated with the TEV-derived C-terminal fragment (lane 7), confirming the fragment mass (lane 7).

^1508_TEV^hNav1.5α^mGFP^, carrying an ENLYFQ TEV cleavage site at residue 1508, migrated in SDS-PAGE also as a ∼250 kDa protein (Fig. 7, lane 1). Co-expression of TEV-encoding cRNA or incubation with recombinant TEV protease after digitonin extraction generated a mGFP-labeled fragment whose mass corresponded to the predicted C-terminal fragment downstream of the TEV site (Fig. 7, lanes 2–3), indicating that both TEV delivery modes, cRNA co-expression and cleavage by commercial TEV are comparably effective. ^SpyT-TEV^hNav1.5α^mGFP-SpyC^, which is capable of SpyT–SpyC-mediated intermolecular ligation, migrated as an apparent ∼500 kDa covalent homodimer that was re-cleaved by TEV into monomeric ∼250 kDa fragments (Fig. 7, lanes 4–5).

Co-expression of ^SpyT-TEV^hNav1.5α^mGFP-SpyC^ with ^SpyT^hNav1.5α^mGFP-SpyT^ further increased homodimerization relative to cleavage (Fig. 7, lanes 6–7), most likely because preferential self-ligation of both constructs, rather than hetero-ligation, generated TEV-less and thus TEV-resistant ^SpyT^hNav1.5α^mGFP-SpyT^ homodimers. A recombinant fragment comprising ^1502–2015^hNav1.5α^mGFP^ comigrated with the C-terminal TEV-derived fragment (Fig. 7, lane 8), confirming the fragment mass. Together, these cleavage and self-ligation patterns support a model in which ^SpyT-TEV^hNav1.5α^mGFP-SpyC^ forms an antiparallel cyclic homodimer that becomes refractory to further labeling by ^SpyC^mGFP once both termini are ligated.

### AlphaFold2 analysis of hNav1.5α supports a stable monomer but no well-defined homodimer

Stimulated by our positive experience with AF2-modeled hP2X7R structures as a basis for interpreting the structural impact of hP2X7R mutations (see next chapter), we modeled both monomeric and homodimeric assemblies of hNav1.5α using AlphaFold2-Multimer (multimer_v3) as implemented in ColabFold, providing the full-length hNav1.5α sequence once or twice (hNav1.5α:hNav1.5α). Multiple sequence alignments were generated automatically with MMseqs2 under default settings; for each run, we sampled multiple random seeds with up to 12 recycles per model, followed by AMBER relaxation. Models were ranked according to the combined AF2-Multimer confidence score (ipTM × 0.8 + pTM × 0.2). Single-chain predictions converged on a moderately confident monomeric fold, with mean pLDDT values of ∼67 and pTM scores of up to 0.70, and additional sampling did not substantially improving these metrics (Fig. 8 A). In contrast, even the top-ranked homodimer predictions showed only moderate overall confidence (pLDDT ∼55–60) and low interface confidence (ipTM ∼0.27–0.30, pTM ∼0.48–0.49), with one subunit adopting a compact, monomer-like architecture while the second chain remained largely disordered (Fig. 8 B). No models with high-confidence, symmetry-consistent homodimer interfaces (ipTM ≥ 0.5) were obtained under any condition tested.

**Figure 8.**
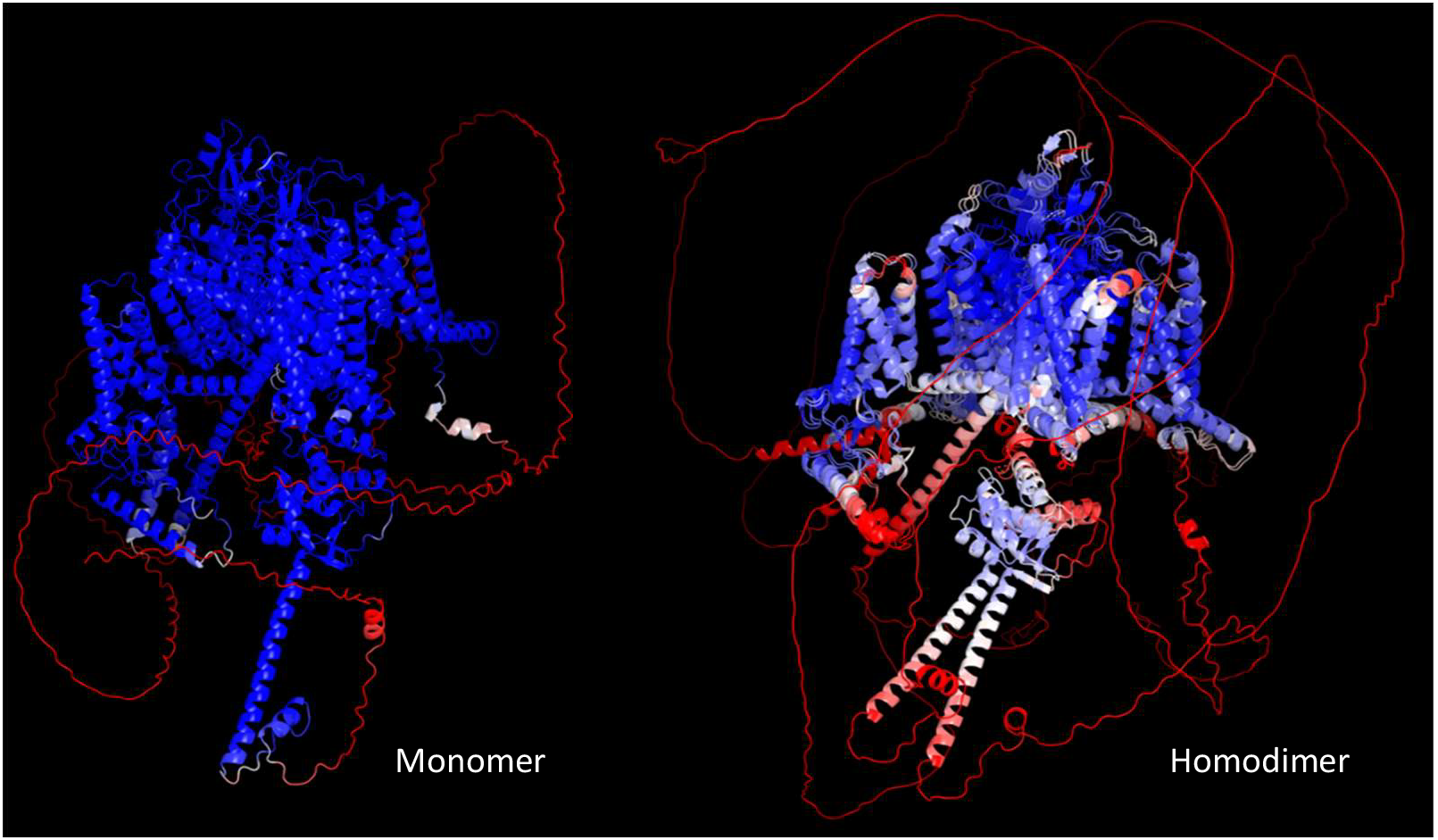
AlphaFold2-Multimer predictions of monomeric homodimer hNav1.5α. Left (**A**), AF2-Multimer (multimer_v3) model of monomeric hNav1.5α, showing a compact helical Nav1.5 core with high local confidence (helices predominantly blue) and extended, low-confidence peripheral regions (red coils) when coloured by pLDDT from red (low) to blue (high). (**B**) AF2-Multimer prediction of the hNav1.5α homodimer, displayed in the same orientation and pLDDT colour scale, in which one hNav1.5α subunit adopts a monomer-like, high-confidence core while the second hNav1.5α subunit is largely disordered and contributes an ill-defined interface (overall pLDDT ∼55–60, ipTM ∼0.27–0.30, pTM ∼0.48–0.49). Both models were generated with ColabFold (AlphaFold2-Multimer multimer_v3, MMseqs2 MSAs, multiple random seeds and recycles, B-factors mapped to pLDDT) and rendered in PyMOL with identical camera settings, emphasizing that AF2 robustly supports a stable monomeric architecture for hNav1.5α but does not identify a high-confidence homodimeric assembly under the conditions used.

Together, these results indicate that AF2 reliably predicts a stable hNav1.5α monomer but fails to generate a consistent, well-defined homodimer model, suggesting that the hNav1.5α–hNav1.5α interface is not strongly sequence-encoded and should be inferred primarily from our experimental data (see Figs. 1-7). The intracellular hNav1.5α homodimers detected under native extraction conditions may therefore reflect non-productive assemblies retained by ER quality control mechanisms a notion supported by the distorted folding of the second protomer in our homodimer predictions or, alternatively, a regulated intracellular reservoir competent for stimulus-dependent forward trafficking to the plasma membrane.

## Discussion

Our combined biochemical and computational analysis resolves a longstanding discrepancy between electrophysiological and structural studies, which have consistently described hNav1.5α as a monomeric channel (Morris & Juranka, 2007; Biswas et al., 2025; Jiang et al., 2021; Li et al., 2021; Pan et al., 2021; Pan et al., 2018; Shen et al., 2017), and biochemical reports of Nav1.5α homodimers with coupled gating (Clatot et al., 2012; Clatot et al., 2017; Rühlmann et al., 2020). By separately tracking the total and plasma-membrane-resident pools of hNav1.5α, we show that both descriptions are correct but refer to distinct subcellular populations: an intracellular, homodimeric pool bearing immature N-glycans, and a plasma-membrane pool that is exclusively monomeric and carries mature, Golgi-processed glycans. This spatial segregation, rather than a true structural discrepancy, likely explains why cryo-EM structures — typically obtained from size-exclusion-purified, monodisperse channel preparations derived from total membrane extracts rather than from a plasma-membrane-selective source — have preferentially captured a monomeric species, whereas biochemical pulldowns of total cellular protein, which do not distinguish subcellular origin, have repeatedly detected the dimer.

What is the physiological role, if any, of the intracellular hNav1.5α homodimer? Several observations argue against it being a simple biosynthetic intermediate en route to the plasma membrane. First, its persistence as a discrete, LiDS-sensitive species independent of the identity or presence of co-expressed β subunits (Fig. 2, Fig. 3) indicates that dimerization is an intrinsic property of the α subunit itself rather than a β-subunit-dependent assembly step. Second, the Endo H/PNGase F sensitivity of the intracellular pool (Fig. 4) places it in the ER or early secretory pathway, compartments subject to stringent quality-control surveillance of membrane protein folding and oligomeric state. We therefore favor a model in which the antiparallel homodimer represents either (i) a non-productive assembly retained by ER quality control, consistent with the poorly defined AlphaFold2-Multimer interface (Fig. 8B), or (ii) a regulated intracellular reservoir that could, in principle, be mobilized for stimulus-dependent forward trafficking. Distinguishing between these possibilities will require pulse-chase experiments tracking the fate of the dimeric pool over time, ideally combined with manipulation of ER-associated degradation (ERAD) components or ER-exit signals.

A central finding of our study is that the intracellular hNav1.5α homodimer adopts a defined antiparallel, cyclic topology, established through the refractoriness of the double-tagged ^SpyC^hNav1.5α^SpyT^ construct to intermolecular labeling (Fig. 6) and confirmed by TEV-cleavage mapping (Fig. 7). This arrangement is an intrinsic property of the channel rather than a consequence of the SpyTag/SpyCatcher design: efficient ligation of the N-terminal SpyC and C-terminal SpyT moieties requires that the N-terminus of one subunit be brought into close proximity to the C-terminus of the neighboring subunit, so that the cyclic ligation product and its TEV-cleavage pattern selectively report a pre-existing topology. To our knowledge, this is the first direct topological assignment of a Nav1.5α– Nav1.5α interface. Conceptually, our strategy parallels the biochemical and structural work that first established the homodimeric architecture of the calcium-activated chloride channel TMEM16A/anoctamin-1 (Fallah et al., 2011; Stolz et al., 2015; Tien et al., 2013; Paulino et al., 2017): in both cases, native electrophoresis and engineered cross-linking or ligation strategies preceded, and predicted features later confirmed by, high-resolution structural methods. We emphasize, however, that the antiparallel arrangement we describe for hNav1.5α is topologically distinct from, and not simply analogous in molecular detail to, the dimer interface of TMEM16A; the comparison is offered at the level of methodological strategy and general precedent for oligomeric ion channel biogenesis, not of shared structural mechanism. Interestingly, a crystal structure of the isolated Nav1.5α C-terminal cytoplasmic domain in complex with calmodulin reported an asymmetric Nav1.5α–Nav1.5α contact mediated by the EF-hand-like domain and helix αVI, with several arrhythmia-associated mutations mapping to this interface (Gabelli et al., 2014). While this soluble-domain contact does not by itself establish the topology of a full-length channel dimer, it raises the possibility that the antiparallel, cyclic dimer we describe biochemically is nucleated, at least in part, by cytoplasmic C-terminal contacts of the kind captured in that crystal structure, rather than solely by the DI–DII linker/14-3-3 interaction proposed by Clatot et al. (2012, 2017).

Our findings also bear on the proposed mechanism of coupled gating between Nav1.5α protomers, in which direct α–α interaction via the intracellular DI–DII linker, reinforced by 14-3-3 binding, was suggested to underlie dominant-negative effects of SCN5A N-terminal mutations linked to inherited arrhythmia syndromes (Clatot et al., 2012; Clatot et al., 2017). If, as our data suggest, the biochemically detectable Nav1.5α homodimer is predominantly an intracellular species, then the coupled-gating phenotypes previously attributed to α–α interaction may instead reflect an indirect mechanism — for example, dominant-negative retention of wild-type subunits in ER-resident heterodimers with mutant protomers, reducing the pool of channels reaching the surface, rather than a direct electromechanical coupling between two co-resident surface channels. This reinterpretation remains compatible with dominant-negative disease mechanisms but shifts the proposed locus of pathogenic action from the plasma membrane to the early secretory pathway. Notably, dimer-dependent phenotypes are not restricted to Nav1.5α: uncoupling of dimeric assembly has been shown to restore the phenotype of a pain-linked Nav1.7 channel mutation (Ruhlmann et al., 2020), suggesting that intracellular homodimerization, and its disruption by disease-associated mutations, may be a conserved feature across Nav channel isoforms.

Our study has several limitations. All experiments were performed in the Xenopus laevis oocyte heterologous expression system, which, while well suited for quantitative, side-by-side comparison of total and surface channel pools, differs from native cardiomyocytes in lipid composition, chaperone repertoire, and the presence of endogenous Nav1.5α-interacting partners. We cannot exclude that the balance between monomeric and dimeric hNav1.5α, or the efficiency of ER export, differs in cardiac tissue. Furthermore, although the S3-, S4-, GFP-, and SpyCatcher/SpyTag-tagged hNav1.5α constructs used throughout this study behaved consistently with one another, we cannot formally exclude subtle effects of these tags on oligomerization propensity, although the reproducibility of the antiparallel dimer across multiple independent tagging strategies (Figs. 5–7) argues against a tag-specific artifact. Finally, the AlphaFold2-Multimer analysis, while informative in showing that the antiparallel interface is not strongly sequence-encoded, is inherently limited by the training data available for membrane protein complexes and cannot substitute for an experimentally determined structure of the intracellular dimer.

Future work should aim to resolve the structure of the intracellular hNav1.5α homodimer directly, for example by cryo-EM of ER-enriched membrane fractions or of the covalently locked SpyC/SpyTag-ligated dimer described here, which may be more amenable to structural analysis than the native, non-covalent assembly. It will also be important to test whether disease-associated mutations in the DI–DII linker and other candidate dimerization interfaces alter the efficiency of ER export of the antiparallel dimer, linking the biochemical phenomenon described here to the cellular pathophysiology of Nav1.5α-linked arrhythmia syndromes. More broadly, our study illustrates that a channel subunit’s oligomeric state can be maturation-stage-specific, and highlights native electrophoresis combined with compartment-selective labeling as a general strategy for detecting such stage-specific assemblies that would otherwise be masked in bulk analyses of total cellular protein.

